# Artificially induced torpor during pregnancy impairs fetal growth in mice

**DOI:** 10.1101/2025.07.31.668002

**Authors:** Mitsue Hagihara, Takeshi Sakurai, Kazunari Miyamichi, Kengo Inada

**Author notes:** Correspondence (K.M.), (K.I.).

## Abstract

Fetal development in endothermic mammals relies on a tightly regulated maternal body temperature. In animal models, deviations from normothermia during gestation cause fetal developmental abnormalities. However, the direct and physiological manipulation of the maternal core temperature has been technologically challenging, as traditional methods involve the imposition of external thermal stress. Recent advances in the identification of thermoregulatory neurons in the mouse preoptic area have allowed precise control of maternal body temperature without altering environmental conditions. Here we show that the activation of excitatory neurons in the medial preoptic area (MPO) induces a torpor-like hypothermic state in pregnant mice. When induced during early gestation, this state resulted in pregnancy loss, likely due to implantation failure. Hypothermia in mid- and late-gestation reduced fetal liver and kidney size and caused severe growth retardation. Similar outcomes were observed following pyroglutamylated RFamide peptide (Qrfp)-expressing neuron activation in the MPO, whereas inhibitory neuron activation had minimal effect. Furthermore, activating MPO excitatory neurons projecting to the dorsomedial hypothalamic nucleus reproduced both the torpor-like state and fetal growth retardation. These results underscore the importance of a stable maternal body temperature in fetal development and establish a new model for studying the effects of altered maternal temperature on embryogenesis.

## Introduction

The physiological state of the maternal body is crucial for fetal development^1^. Among the various factors, maternal body temperature, which is regulated by the brain, is particularly critical. Deviations from normothermia is known to adversely affect fetuses. Human clinical studies have shown that maternal body temperature is closely correlated with fetal heart rate, and maternal hypothermic conditions lead to fetal bradycardia^2-4^. In such cases, emergency rewarming procedures are important to prevent miscarriages or developmental anomalies^2,4,5^. Animal model studies are essential for deciphering the detailed developmental effects of altered thermal conditions. A classical study in dogs revealed that maternal hypothermia decreases uterine blood flow^6^, implying that fetal nutrient and oxygen supply may be compromised under hypothermic stress. In rodent studies, maternal thermal manipulation is typically achieved through environmental temperature shifts by exposing pregnant females to hot or cold conditions. These studies have revealed that both hyperthermic and hypothermic environments increase the risk of fetal growth retardation, malformations, and lethality^7-11^. However, these manipulations with extreme thermal challenges confound the interpretation of whether the observed fetal outcomes are direct consequences of thermal/metabolic disturbances.

Studies in rodents have identified the preoptic area of the hypothalamus (POA) as a crucial regulator of body temperature homeostasis^12,13^. Recent studies have further elucidated the molecular underpinnings of various thermoregulatory cell types in this region^14-21^. For instance, warmth-sensitive POA neurons co-expressing pituitary adenylate cyclase-activating polypeptide (PACAP) and brain-derived neurotrophic factor (BDNF) can induce hypothermia upon activation^16^, while prostaglandin E type 3 receptor (EP3R)-expressing neurons mediate febrile responses^21^. Moreover, POA neurons expressing violet light-sensitive opsin 5 (Opn5)^22^, pyroglutamylated RFamide peptide (QRFP)^14^, leptin receptor^15^, transient receptor potential cation channel subfamily M member 2 (Trpm2)^23^, or POA neurons activated during daily torpor^19^ can induce a prolonged torpor-like hypothermic state with temporal precision. Notably, many of these thermoregulatory neurons co-express vesicular glutamate transporter type 2 (vGluT2)^24^, which is consistent with the findings that vGluT2-expressing (vGluT2+) neurons can also drive a torpor-like state^15^. Some POA neurons are thought to project to downstream neural circuits, including pathways extending from the dorsomedial hypothalamic nucleus (DMH) to the medullary raphe nuclei, eventually controlling the sympathetic thermoregulatory systems^12,25^.

These recent advances in our understanding of thermoregulatory centers within the POA have made it possible to modulate maternal body temperature with temporal precision without the need for extreme thermal challenges. In this study, we leveraged this mechanistic framework to investigate how the induction of a torpor-like hypothermic state in pregnant mice affects fetal growth. Using the chemogenetic activation of excitatory neurons in the medial preoptic area (MPO) defined in the Allen Brain Atlas^26^, which is abbreviated as MPA in the Paxinos atlas^27^, we demonstrate that maternal hypothermia markedly impairs fetal growth and organ development.

## Results

### Maternal torpor-like state during early gestation terminates pregnancy

We first confirmed that a torpor-like hypothermic state can be induced at various stages of pregnancy using the chemogenetic activation of MPO excitatory neurons^15,28^. An adeno-associated virus (AAV) encoding Cre-dependent hM3Dq was injected into the MPO of *vGluT2-Cre* female mice (Fig. 1a and 1b). Three weeks after the viral injection, these female mice were mated with male mice. The day of confirmation of the vaginal plug was designated as gestational day (GD) 0.5. We administered intraperitoneal injections of clozapine N-oxide (CNO) or saline on GD 5.5, 9.5, 12.5, or 15.5, and measured maternal rectal temperature for the following three days (Fig. 1c). Consistent with previous studies^15,20^, CNO injection decreased the rectal temperature to below 30 °C within several hours, which was not observed in the saline-injected controls (Fig. 1d). In most mice of the CNO-injected groups, rectal temperature recovered to above 30 °C within 72 h. However, CNO injection on GD 15.5 exhibited a prolonged hypothermic state. Although the minimum rectal temperature reached during the 72-h period was comparable across gestational stages (Fig. 1e), the rectal temperature remained significantly lower at 72 h in the GD 15.5 CNO group (Fig. 1f), suggesting impaired thermoregulatory recovery in late pregnancy.

**Fig. 1.**
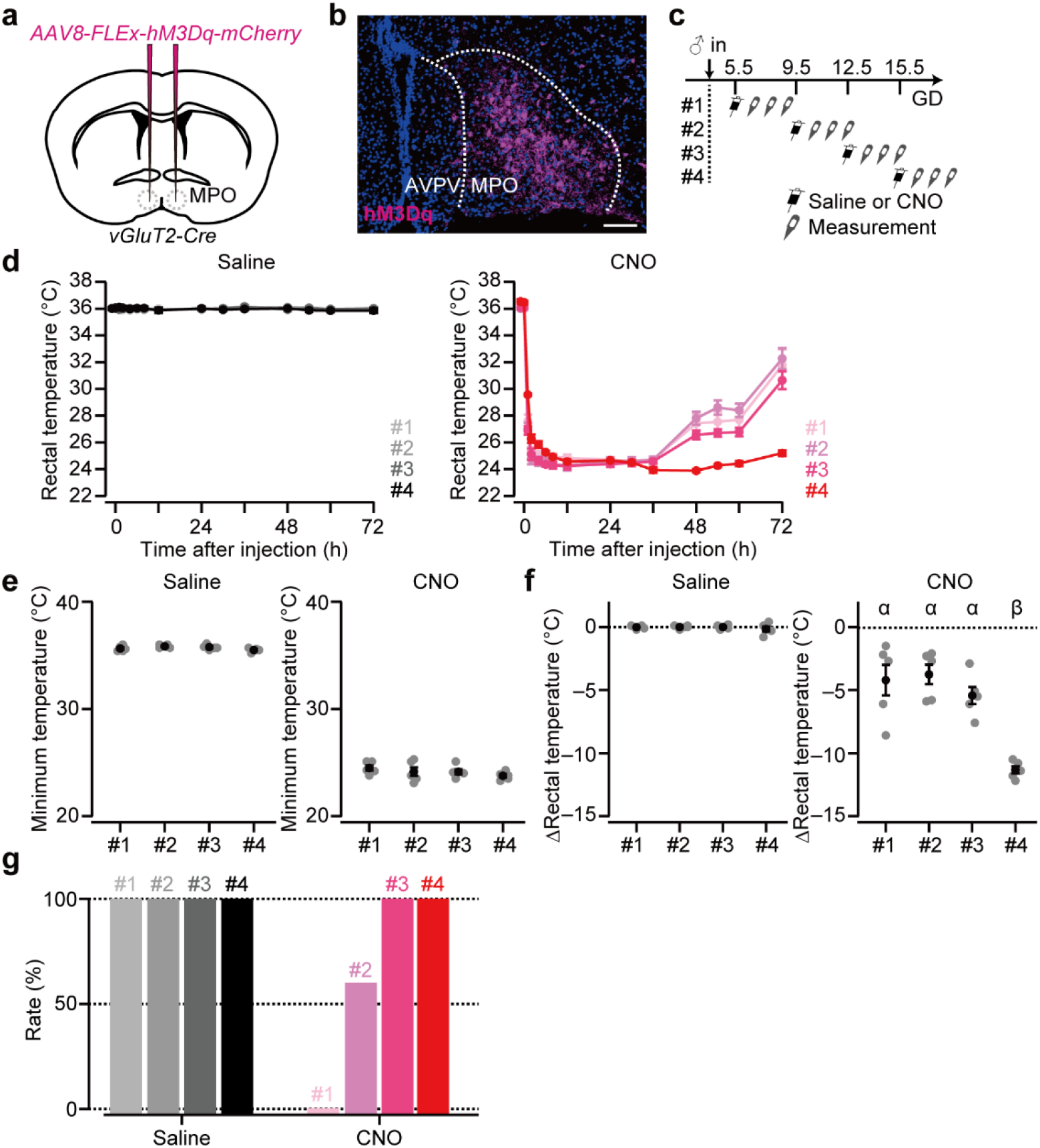
Chemogenetic activation of excitatory neurons in the MPO induces a torpor-like hypothermic state in pregnant female mice. **(a)** Schematic of the virus injection. *AAV-FLEx-hM3Dq-mCherry* is injected bilaterally into the MPO of *vGluT2-Cre* mice. (b) Representative coronal section of the MPO showing hM3Dq-mCherry expression (magenta). Blue, DAPI. Scale bar, 50 µm. (**c**) Schematic of the experimental timeline. Intraperitoneal injections are administered on gestation day (GD) 5.5, 9.5, 12.5, or 15.5, referred to as timepoints #1, #2, #3, and #4, respectively. Rectal temperature is measured for three consecutive days after the intraperitoneal injection. (**d**) Time course of the rectal temperature of pregnant females after saline (left) or CNO (right) injections. Note that intraperitoneal injection is performed at zeitgeber time 0. n = 5 mice each. (**e**) Minimum rectal temperature within 72 h after saline or CNO injection. We did not find statistical significance in both groups (saline, p = 0.094, CNO, p = 0.41, one-way ANOVA). (**f**) Change in rectal temperature, calculated as the difference between 72 h after (T_72_) and 1 h before (T_−1_) the intraperitoneal injection of saline or CNO. Different letters (α and β) in the upper part of the graph denote significant differences at p < 0.01 by one-way ANOVA with post-hoc Tukey HSD (#1 versus #4, p = 0.0010, #2 versus #4, p = 0.0010, #3 versus #4, p = 0.0014). (**g**) Percentage of female mice that maintained fetuses three days after saline or CNO injection. Note, this phenotype exhibited an all-or-none pattern: each dam either retained the entire litter or experienced complete fetal loss. Color coding corresponds to panel **d**. n = 5 mice each. Error bars, standard error of the mean.

To assess the impact of this torpor-like state on pregnancy continuation, we performed Caesarean sections three days after CNO injection. Fetuses were present in all mice treated with CNO on GD 12.5 and 15.5. In contrast, a fraction of dams experienced fetal loss in the GD 9.5 CNO group (#2 in Fig. 1g), and no fetuses or implantation scars were found in the GD 5.5 CNO group (#1 in Fig. 1g) despite prior mating, as indicated by vaginal plugs. Saline-injected controls maintained normal pregnancies at all the time points. These results demonstrate that the induction of a torpor-like state during early gestation severely compromises pregnancy, likely by disrupting implantation processes and/or subsequent early embryonic development.

### Maternal torpor-like state delays fetal organ growth

To analyze the influence of the maternal torpor-like hypothermic state on fetal development, we conducted histological analyses of the fetuses. To achieve this, we prepared pregnant female mice expressing hM3Dq in the *vGluT2+* neurons in the MPO, and injected CNO or saline on GD 12.5, a time when organs such as the heart, liver, and kidney undergo rapid growth^29-31^. The fetuses were collected via Caesarean section on GD 15.5 (Fig. 2a). Fetal body weight was significantly lower in the CNO-injected group than in the saline-injected group, demonstrating that the maternal torpor-like state severely impaired overall fetal growth (Fig. 2b and 2c). From each litter, one fetus was randomly selected, and two sagittal sections were prepared: one containing the lateral ventricle of the brain and the center of the heart and liver, and the other containing the kidney. Sections were stained with hematoxylin and eosin (HE) (Fig. 2d). The organ areas in these sections were quantified. Fetuses from hypothermic dams showed significantly smaller liver and kidney sizes than normothermic mice, whereas the heart size remained comparable (Fig. 2e). The lateral ventricle of the brain appeared enlarged in the CNO group, although this trend did not reach statistical significance (p = 0.066, two-sided Mann–Whitney *U*-test). These results suggest that the maternal torpor-like state on GD 12.5–15.5 disturbs fetal growth in an organ-specific manner.

**Fig. 2.**
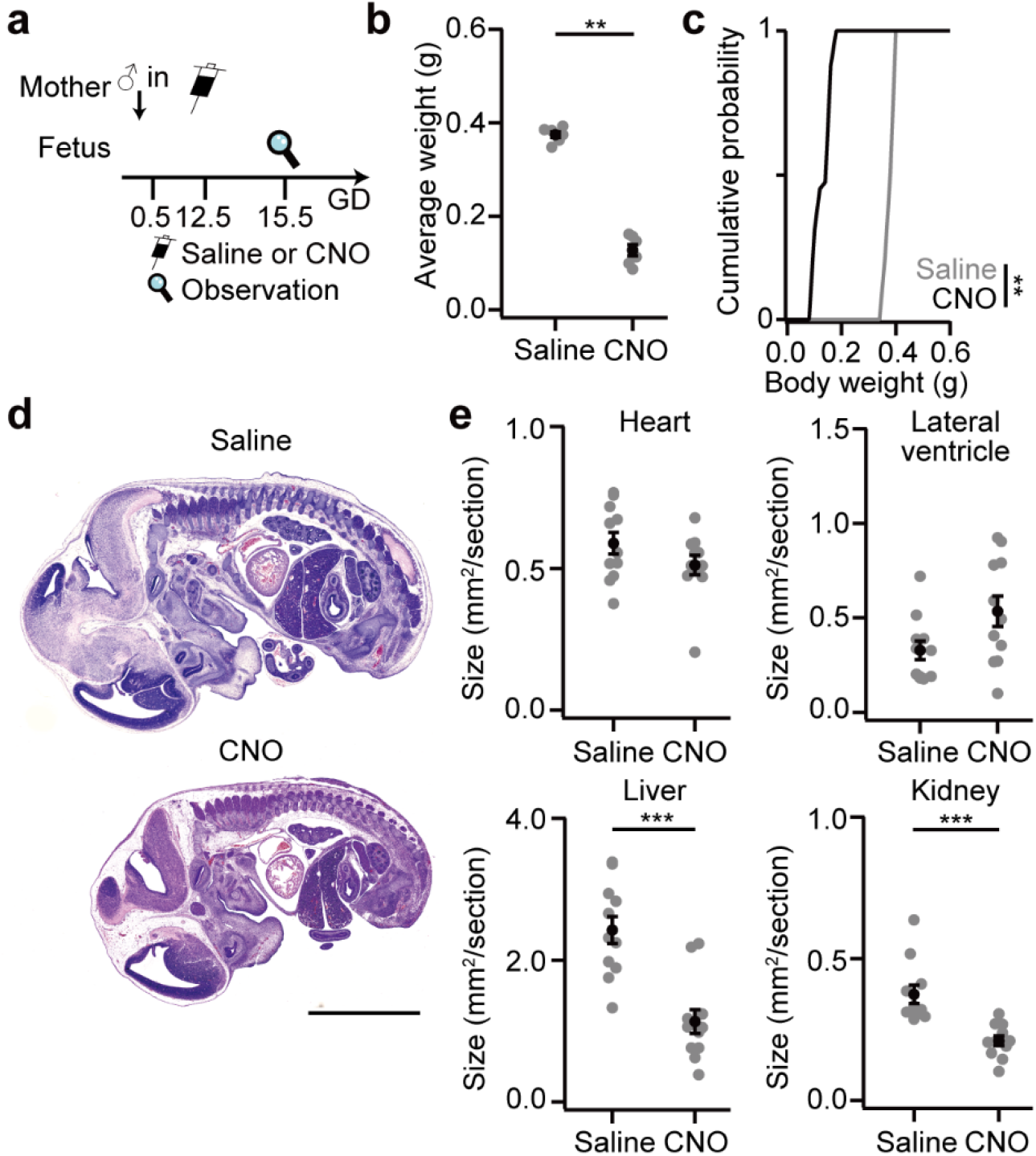
The maternal hypothermic state delays fetal organ growth in a tissue specific manner. (**a**) Schematic of the experimental timeline. Saline or CNO is injected into the pregnant female mice expressing hM3Dq in the MPO *vGluT2+* neurons on GD 12.5. (**b**) Average fetal body weight per dam (**p = 0.0051, two-sided Mann–Whitney *U*-test, n = 6 dams each). (**c**) Cumulative probability of fetal body weights (**p = 0.0035, Kolmogorov–Smirnov test). Saline, n = 47 fetuses from 6 dams; CNO, n = 42 fetuses from 6 dams. (**d**) Representative images of fetuses collected on GD 15.5 from the dams injected with saline (top) or CNO (bottom) on GD 12.5. Scale bar, 300 µm. (**e**) Size of the heart, lateral ventricle in the brain, liver, and kidney (***p < 0.001, two-sided Mann–Whitney *U*-test. The p values are: heart, p = 0.29; lateral ventricle, p = 0.066; liver, p = 0.0006; kidney, p = 0.0001. n = 11 fetuses each). Error bars, standard error of the mean.

### Maternal torpor-like state during late pregnancy causes fetal lethality

To unveil the impacts of maternal hypothermia during late gestation, we chemogenetically activated MPO excitatory neurons on GD 15.5 and performed Caesarean sections on GD 16.5 to 18.5 (Fig. 3a and 3b). Fetal viability was determined based on the presence of blood flow or spontaneous movement under stereomicroscopy. All fetuses from the saline-injected control dams were alive at all time points (Fig. 3c). Similarly, most fetuses from the CNO-injected dams remained viable on GD 16.5 (Fig. 3c), indicating that a single day of maternal hypothermia does not cause immediate lethality. These data are consistent with those of a previous study showing that short-term maternal hypothermia does not compromise pregnancy maintenance in mice^28^. However, the fetal survival rate declined by GD 17.5, and the majority of fetuses were nonviable by GD 18.5 (Fig. 3c), indicating that prolonged maternal hypothermia is deleterious during late gestation. Fetuses from the CNO-injected dams showed significantly reduced body weights compared to fetuses from the saline-injected dams, even at GD 16.5 and GD 17.5 (Fig. 3d). The fetal weight distribution in the CNO group differed substantially from that in the saline controls (Fig. 3e). Collectively, these results suggest that a sustained maternal hypothermic state induced by the activation of MPO excitatory neurons impairs fetal growth and ultimately results in lethality. This fetal demise may have contributed to the prolonged hypothermic state observed during late gestation (Fig. 1d), possibly due to disrupted maternal health. Alternatively, altered maternal metabolism during late pregnancy leading to delayed degradation of CNO, thereby prolonging its thermoregulatory effects is also a possibility.

**Fig. 3.**
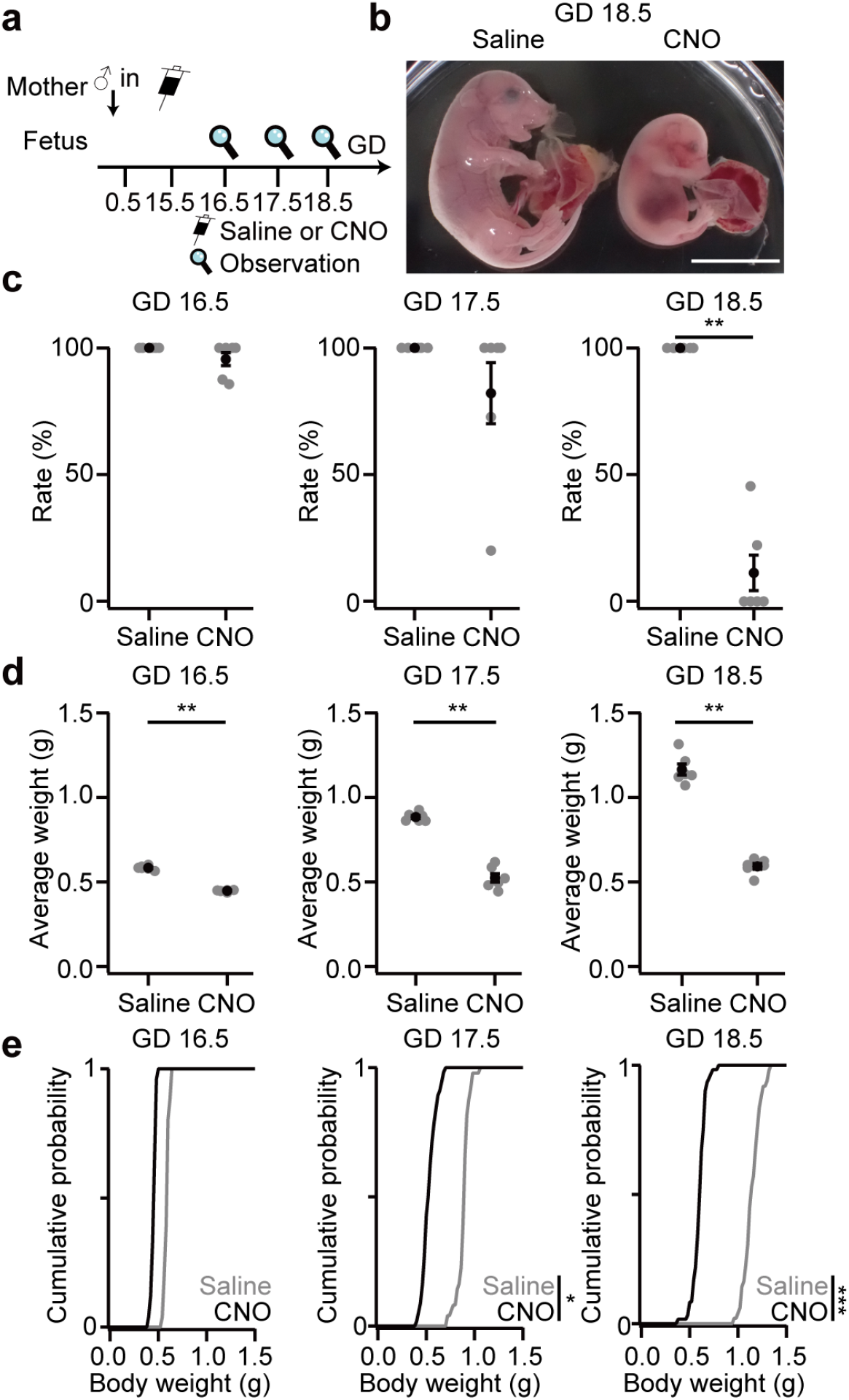
The maternal hypothermic state impairs fetal weight gain. (**a**) Schematic of the experimental timeline. Saline or CNO was intraperitoneally injected into pregnant female mice expressing hM3Dq in the *vGluT2+* MPO neurons on gestation day (GD) 15.5. Caesarean sections are performed on GD 16.5, 17.5, and 18.5. (**b**) Representative image of fetuses from GD 18.5 dams injected with saline (left) or CNO injection (right) on GD 15.5. Scale bar, 1 cm. (**c**) Survival rate of fetuses per dam (**p < 0.01, two-sided Mann–Whitney *U*-test. The p values are: GD 16.5, p = 0.38, GD 17.5, p = 0.38, GD 18.5, p = 0.0051. n = 6 mothers each). (**d**) Average fetal body weight per dam (**p = 0.0051, two-sided Mann–Whitney *U*-test). (**e**) Cumulative probability of fetal body weights (*p = 0.010, ***p = 6.4 × 10^−6^, Kolmogorov– Smirnov test). n = 49–60 fetuses from 6 dams each. Error bars, standard error of the mean.

As a control, the same experimental procedure was repeated using pregnant females expressing mCherry alone. We found that intraperitoneal injection of CNO on GD 15.5 neither evoked hypothermia (Supplementary Fig. S1a–1e) nor affected fetal viability or body weight (Supplementary Fig. S1f–1h). Additionally, we chemogenetically activated *vesicular GABA transporter* (*vGAT*)-expressing neurons in the MPO (Supplementary Fig. S1i and 1j), based on a previous study showing that POA-inhibitory neurons do not evoke hypothermia^15^. In this cohort, CNO injection on GD 15.5 led to a modest drop in rectal temperature (Supplementary Fig. S1k and 1l), but values remained above 30 °C, and normothermia was restored within 24 h (Supplementary Fig. S1k–1m). Fetal body weights did not differ significantly between the CNO- and saline-injected groups (Supplementary Fig. S1n–1p). These results confirmed that hypothermia-induced fetal growth impairment and lethality are specific to the activation of excitatory *vGluT2+* neurons in the MPO.

### *Qrfp*-expressing neurons recapitulate the phenotypes observed in *vGluT2-Cre* mice

A recent study showed that *Qrfp*-expressing (*Qrfp+*) neurons in the MPO can induce a hibernation-like state termed Q-induced hypometabolism (QIH)^14^. Given that most *Qrfp+* neurons co-express *vGluT2*^14^, we hypothesized that the chemogenetic activation of MPO *Qrfp+* neurons reproduces the phenotypes observed in *vGluT2-Cre* mice. To test this, we used *Qrfp-iCre* mice expressing hM3Dq in MPO neurons (Fig. 4a and 4b) and injected either CNO or saline on GD 15.5 (Fig. 4c). Similar to that previously reported^14^, CNO treatment reliably induced QIH, as shown by the decrease in body temperature (Fig. 4d–4f). This state mirrored the physiological state in *vGluT2-Cre* mice; CNO-injected dams exhibited significantly lower minimum rectal temperatures and prolonged hypothermia at 72 h compared to saline-injected controls (Fig. 4d–4f). Moreover, by GD 18.5, the fetal survival rate significantly reduced (Fig. 4g), with significantly lower fetal body weights compared with the controls (Fig 4h and 4i). These results suggest that the chemogenetic activation of *Qrfp+* neurons in the MPO largely recapitulates the hypothermic and developmental phenotypes observed with the activation of *vGluT2+* neurons.

**Fig. 4.**
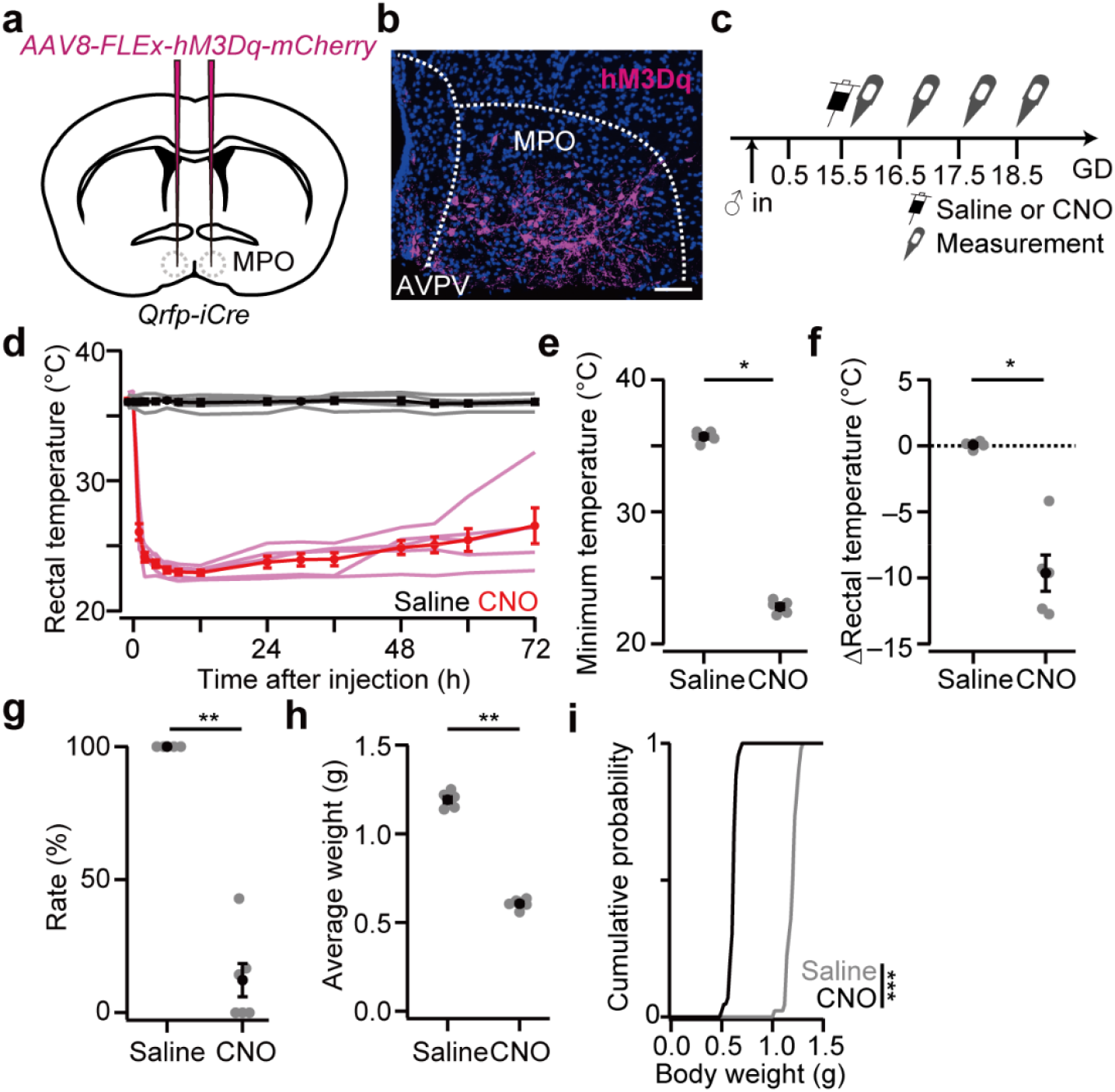
*Qrfp*-expressing neurons largely recapitulate phenotypes observed in *vGluT2*-expressing neurons. (**a**) Schematic of the virus injection. *AAV-FLEx-hM3Dq-mCherry* injected bilaterally into the MPO of *Qrfp-iCre* mice. (**b**) Representative coronal section of the MPO showing hM3Dq-mCherry expression (magenta). Blue, DAPI. AVPV, anteroventral periventricular nucleus. Scale bar, 50 µm. (**c**) Schematic of the experimental timeline. (**d**) Time course of rectal temperature of pregnant female mice after intraperitoneal injection of saline (black) or CNO (magenta) on GD 15.5. n = 5 mice each. (**e**) Minimum rectal temperature within the 72-h window after injection (*p = 0.012, two-sided Mann–Whitney *U*-test). (**f**) Change in rectal temperature (T_72_ − T_−1_) (*p = 0.012, two-sided Mann–Whitney *U*-test). (**g**) Survival rate of fetuses per dam (**p = 0.0051, two-sided Mann–Whitney *U*-test. n = 6 mothers each). (**h**) Average fetal body weight per dam (**p = 0.0051, two-sided Mann–Whitney *U*-test). (**i**) Cumulative probability of fetal body weight (***p = 6.4 × 10^−6^, Kolmogorov–Smirnov test). Saline, n = 45 fetuses from 6 dams, CNO, n = 43 fetuses from 6 dams. Error bars, standard error of the mean.

### MPO excitatory projections to DMH contribute to maternal hypothermia and fetal phenotypes

To characterize the downstream pathways by which MPO excitatory neurons induce hypothermic states during pregnancy, we focused on the DMH, a key node in thermoregulatory circuits, including those for torpor^12,13,23,25^. A previous study showed that *Qrfp+* neurons projecting to the DMH are responsible for QIH^14^, raising the possibility that MPO *vGluT2+* neurons may act similarly through the DMH to mediate hypothermia in pregnant females. First, we visualized the axons of MPO *vGluT2+* neurons by injecting *AAV-FLEx-ChR2-EYFP* into the MPO of *vGluT2-Cre* mice (Fig. 5a). Immunostaining three weeks after the injection showed that these neurons sent their axons broadly to brain regions, including the arcuate hypothalamic nucleus (ARH), ventromedial hypothalamic nucleus (VMH), and lateral hypothalamic area (LHA) (Fig. 5b). Dense enhanced yellow fluorescent protein (EYFP)-positive fibers were observed in the DMH, indicating intensive MPO-to-DMH projections (Fig. 5b).

**Fig. 5.**
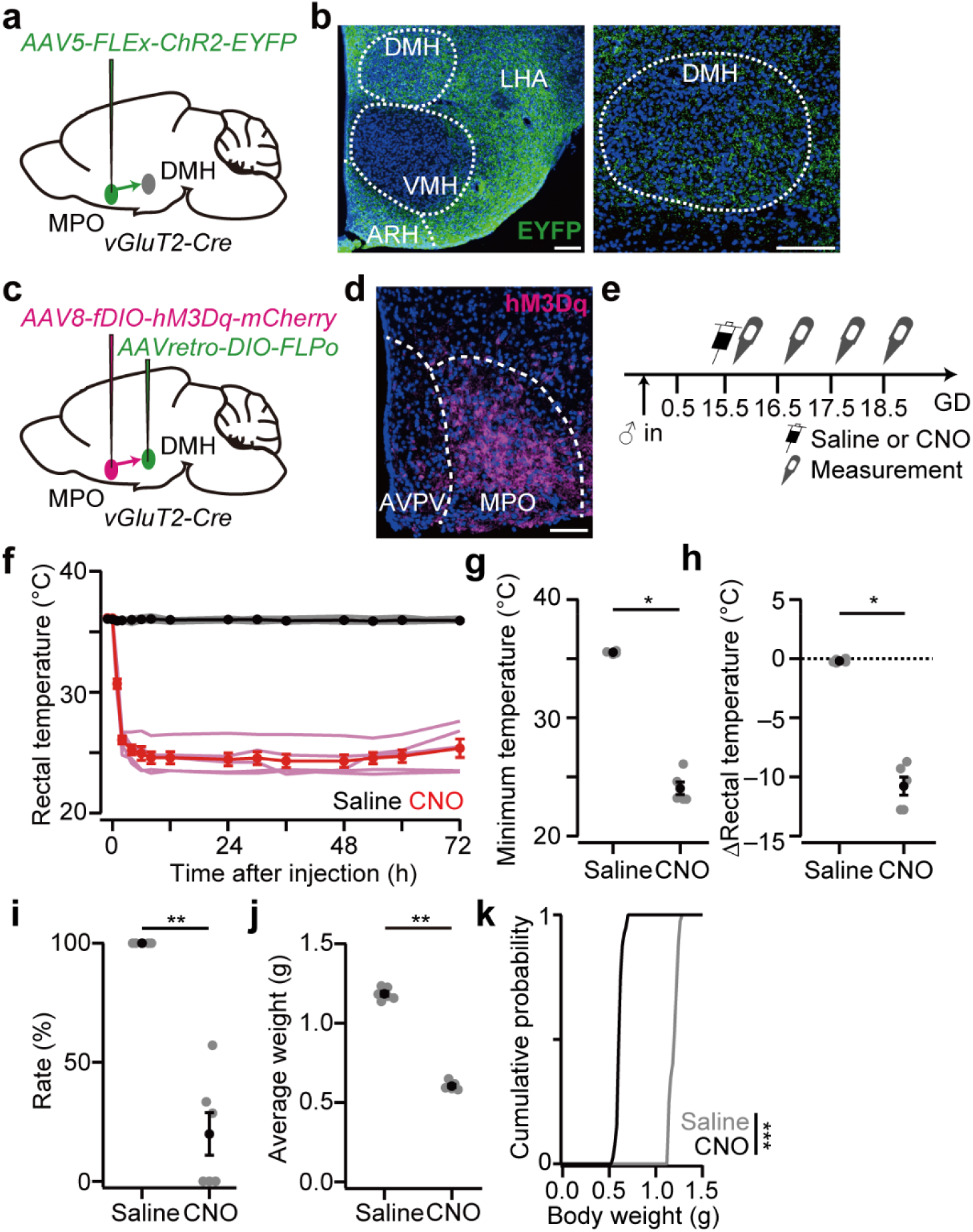
Excitatory projections from MPO to DMH recapitulate the hypothermic state and fetal phenotypes. (**a**) Schematic of AAV injection. *AAV-FLEx-ChR2-EYFP* is injected unilaterally into the MPO of *vGluT2-Cre* mice to visualize the excitatory projections. (**b**) Representative coronal sections showing EYFP-labeled axonal projections (green). Blue, DAPI. Scale bars, 50 µm. (**c**) Schematic of the virus injection. *AAVretro-FLEx-FLPo* is injected bilaterally into the DMH of *vGluT2-Cre* mice, followed by an injection of *AAV-fDIO-hM3Dq-mCherry* into the bilateral MPO. (**d**) Representative coronal section of the MPO showing hM3Dq-mCherry expression (magenta). Blue, DAPI. Scale bar, 50 µm. (**e**) Schematic of the experimental timeline. (**f**) Time course of the rectal temperature after intraperitoneal injection of saline (black) or CNO (magenta) on GD 15.5. n = 5 mice each. (**g**) Minimum rectal temperature within the 72-h window after injection (*p = 0.012, two-sided Mann–Whitney *U*-test). (**h**) Change in rectal temperature (T_72_ − T_−1_) (*p = 0.012, two-sided Mann–Whitney *U*-test). (**i**) Survival rate of fetuses per dam (**p = 0.0051, two-sided Mann–Whitney *U*-test. n = 6 dam each). (**j**) Average fetal body weight per dam (**p = 0.0051, two-sided Mann–Whitney *U*-test). (**k**) Cumulative probability of fetal body weights (***p = 6.4 × 10^−6^, Kolmogorov–Smirnov test). n = 40 fetuses from 6 dams each. Error bars, standard error of the mean.

To test whether this pathway mediates maternal hypothermia and its subsequent effects on fetuses, we selectively expressed hM3Dq in DMH-projecting MPO *vGluT2+* neurons using a retrograde viral strategy (Fig. 5c and 5d). CNO or saline injections were administered to the mice on GD 15.5 (Fig. 5e). Chemogenetic activation of DMH-projecting MPO *vGluT2+* neurons reliably evoked hypothermia (Fig. 5f–5h), significantly decreased fetal survival by GD 18.5 (Figure 5i), and led to lower fetal body weights (Fig. 5j and 5k). Collectively, these results confirm the critical role of excitatory MPO neurons projecting to the DMH in body temperature regulation, and further suggest that this pathway mediates hypothermia-induced impairments in fetal development in pregnant female mice.

## Discussion

Mammals maintain a remarkably stable body temperature, typically higher than ambient conditions, through endothermic mechanisms. This thermo-regulatory stability supports robust embryonic development. Although classical studies have shown that altering the maternal body temperature increases the risk of fetal growth retardation, malformations, and lethality^7-11^, these findings were obtained using extreme thermal challenges. Direct manipulation of the maternal body temperature set-point in a controlled and physiological manner has long been a challenge. To this end, we introduced recently established methods to induce a torpor-like hypothermic state via the chemogenetic activation of specific neural populations in the MPO of pregnant female mice. While such torpor-like states in mice have been suggested to have anti-aging effects and therapeutic benefits in disease models^32,33^, its impact on reproductive function remains largely uncharacterized. This approach enabled the direct and temporally precise manipulation of maternal body temperature, allowing us to assess its causal impact on pregnancy outcomes.

Our data demonstrated that torpor-like hypothermic states can be induced in pregnant female mice with an efficiency comparable to that observed in non-pregnant females and males in previous studies^14,15^. When hypothermia was induced at an early gestational stage (GD 5.5), we observed complete failure of pregnancy continuation, suggesting that a normothermic environment is critical for early pregnancy maintenance (Fig. 1). In contrast, hypothermia at later stages did not lead to miscarriage but severely disturbed fetal growth and organ development (Figs. 2 and 3). These results are consistent with and expand on the classical observation that maternal hypothermia affects pregnancy outcomes in a gestational stage-dependent manner^7-9^.

We found that the maternal torpor-like state initiated on GD 12.5 disturbs fetal organ growth in a tissue-specific manner. Specifically, CNO administration attenuated the growth of the fetal kidney and liver, while sparing the heart (Fig. 2), despite the rapid growth of all three organs around GD 12.5^29-31^. The underlying mechanisms driving this differential sensitivity to maternal hypothermic state remain unclear. One possibility is that fetal organs may experience distinct local temperatures during maternal hypothermia, as reported in a previous study on organ-specific temperature variations^34^. Alternatively, certain organs may have intrinsic mechanisms that respond to fluctuations in the ambient temperature. For example, incubation at a suboptimal ambient temperature induces significant enlargement of the heart but not of the kidneys^35^. Future investigations into the gene expression patterns that regulate cell differentiation and tissue formation in each fetal organ may elucidate the mechanisms underlying the tissue-specific effects of maternal hypothermia.

We also demonstrated that maternal hypothermia on GD 15.5 resulted in a marked attenuation of fetal weight gain between GD 16.5 and 18.5, with weights staying ∼0.5 g (Fig. 3). Given that body temperature affects numerous physiological systems in both the mother and the fetus, this growth retardation likely arises from molecular and/or metabolic disruptions. If fetuses experience a corresponding temperature drop in utero, temperature-sensitive gene expression dynamics may be disturbed. For example, a lower ambient temperature delays the oscillatory expression of *deltaC*, a Notch signaling ligand, in zebrafish embryos^36^. A similar disturbance in mammalian gene expression rhythms can disturb proper embryogenesis and fetal weight gain. However, studies on sheep^37^ and baboons^38^ suggest that fetal body temperature fluctuations are buffered relative to maternal fluctuations. If this is true in our mouse model, impaired maternal energy metabolism^14,19^ or decreased uterine blood flow^6^ may compromise nutrient and oxygen delivery to the fetus, leading to disturbed fetal development. Supporting this view, chick embryos incubated at non-optimal temperatures show increased lethality, malformations, and growth delay^35,39^, underscoring the importance of normothermia in fetal development. Future studies should examine intrafetal thermal distribution and gene expression dynamics to elucidate how hypothermia perturbs specific developmental programs. More generally, our established model offers a useful platform to further investigate the potential influence of torpor-like states on fetal development and the reproductive health of pregnant females.

## Methods

### Ethics declarations

All experimental procedures were approved by the Institutional Animal Care and Use Committee of the RIKEN Kobe Branch.

### Animals

The animals were housed under a 12-h light/12-h dark cycle with *ad libitum* access to food and water. Zeitgeber time 0 corresponds to 8:00 am. Wild-type C57BL/6J mice were purchased from Japan SLC. *vGluT2-Cre* (also known as *Slc17a6-ires-Cre*, Jax #028863) and *vGAT-Cre* (also known as *Slc32a1-ires-Cre*, Jax #028862) were purchased from Jackson Laboratory. *Qrfp-iCre* mouse line^14^ was a kind gift from Dr. Genshiro A. Sunagawa and Dr. Takeshi Sakurai.

### Viral preparations

We obtained the following AAV vectors from Addgene (titer is shown as genome particles [gp] per mL): AAV serotype 8 *hSyn-FLEx-hM3Dq-mCherry* (#44361, 2.1 × 10^13^ gp/mL), AAV serotype 8 *hSyn-FLEx-mCherry* (#50459, 1.7 × 10^13^ gp/mL), AAV serotype 5 *EF1a-FLEx-hChR2(H134R)-EYFP* (#20298, 1.8 × 10^13^ gp/mL), and AAV retrograde *EF1a-DIO-FLPo* (#87306, 2.3 × 10^13^ gp/mL). AAV serotype 8 *hSyn-fDIO-hM3Dq-mCherry* (2.0 × 10^13^ gp/mL) was purchased from the Canadian Neurophotonics Platform Viral Vector Core Facility.

### Stereotactic injection

For targeted AAV injection in a specific brain region, stereotactic coordinates were defined for each brain region using the Allen Mouse Brain Atlas^26^. The mice were anesthetized with 65 mg/kg ketamine (Daiichi Sankyo) and 13 mg/kg xylazine (X1251, Sigma-Aldrich) via intraperitoneal injection and head-fixed to stereotactic equipment (Narishige). The following coordinates were used (in mm from the bregma for anteroposterior [AP], mediolateral [ML], and dorsoventral [DV]): MPO, AP 0.4, ML 0.2, DV 5.2; DMH, AP −1.5, ML 0.2, DV 5.0. The injection volume of the viruses was 200 nL and the speed was 50 nL/min. After the viral injection, the animals were returned to their home cages.

### Measurement of rectal temperature

In Figs. 1–4 and Supplementary Fig. S1, experiments were conducted three weeks after the injection of an AAV expressing *hM3Dq-mCherry* or *mCherry*. In Fig. 5, *AAVretro-DIO-FLPo* was injected bilaterally into the DMH. Two weeks later, *AAV8-fDIO-hM3Dq-mCherry* was injected bilaterally into the MPO. Each female was mated with a wild-type male. After the detection of the vaginal plug, each female mouse was housed individually. The rectal temperature was measured using a thermometer (KN-91-AD1687-M, A&D Company). In Figs.

1d, 4d, and 5f and Supplementary Figs. S1c and 1k, time 0 corresponds to a zeitgeber time of 0. CNO (catalog #4936, Tocris) was dissolved in saline and administered via an intraperitoneal injection at a dose of 1 mg/kg. Because hypothermic animals can exhibit lower locomotor activity^14^, we distributed food pellets and water jelly in the cage. The evidence of chewing confirmed that the animals had access to both food and water.

### Analysis of fetuses

Pregnant female mice were prepared as described above. Each assay was conducted by two experimenters to ensure blinding: Experimenter 1 prepared two identical tubes containing either saline or saline with CNO, and Experimenter 2, who was blinded to the contents, performed the Caesarean section and subsequent analyses. For each fetus, viability was assessed by evaluating blood flow and spontaneous movement detected using a stereomicroscope (SZX 16, Olympus), and fetuses exhibiting either sign were considered alive. Fetal body weight was measured using a precision balance with a resolution of 0.1 mg (ENTRIS224I-1S, Sartorius). Each fetus was fixed in 4% paraformaldehyde (PFA) overnight (Fig. 2). Two four-micron paraffin sections, one containing lateral ventricle of the brain and the center of heart and liver, and the other containing kidney, were prepared. Sections were stained with hematoxylin and eosin (HE) (Genostaff, Tokyo, Japan). Images were acquired using an Olympus BX53 microscope equipped with a 4× (N.A. 0.16) objective lens. Organ area was measured using ImageJ software. The image shown in Fig. 2d was obtained using a slide scanner (AxioScan, Zeiss).

### Histochemistry

The mice were anesthetized with isoflurane and perfused with PBS, followed by 4% paraformaldehyde (PFA) in PBS. The brains were post-fixed overnight in 4% PFA overnight. Twenty-micron coronal brain sections were prepared using a cryostat (Leica). The following primary and secondary antibodies were used for immunostaining at the indicated dilutions: chicken anti-GFP (GFP-1010; Aves Labs; 1:500), rat anti-RFP (5f8; Chromotek; 1:500), anti-chicken Alexa Fluor 488 (703-545-155; Jackson ImmunoResearch; 1:500), and anti-rat Cy3 (712-165-153; Jackson ImmunoResearch; 1:500). Fluoromount (K024; Diagnostic BioSystems) was used as the mounting medium. Brain images were acquired using an Olympus BX53 microscope equipped with a 10× (N.A. 0.4) objective lens.

## Data analysis

All mean values are reported as the mean ± standard error of mean (SEM). The statistical details of each experiment, including the statistical tests used, the exact value of n, and what n represents, are shown in the figure legends. P-values are shown in the figure legends.

## Funding

This work was supported by the JSPS Kakenhi Transformative Research Areas (A) (23H04945, 23H04939) to K.M. and RIKEN Incentive Research Projects to K.I.

## Competing Interests

The authors declare that they have no competing interests.

## Data availability

All data are available in the main paper and supplementary materials.

## Author contributions

K.I. conceived and designed the experiments. M.H. and K.I. performed the experiments and analyzed the data. T.S. provided *Qrfp-iCre* mice. K.M. supervised the study. M. H., K. M., and K. I. discussed the results and drafted the manuscript. K.M. and K.I. edited and reviewed the manuscript.

## Acknowledgements

We thank Drs. Genshiro A. Sunagawa and Takeshi Sakurai for providing transgenic mouse lines. We also thank Wataru Kimura, Teppei Goto, and members of the Miyamichi laboratory for their critical reading of the manuscript.

## Notes

### Competing Interest Statement

The authors have declared no competing interest.

